# Isotopic niches do not follow the expectations of niche conservatism in the bird genus *Cinclodes*

**DOI:** 10.1101/2022.10.10.511651

**Authors:** Jonathan A. Rader, Daniel R. Matute

**Affiliations:** Biology Department, University of North Carolina at Chapel Hill, Chapel Hill, NC 27599, USA

**Keywords:** Stable isotope analysis, morphospace, Furnariidae, trait evolution, macroevolution, phylogenetic analysis

## Abstract

Phenotypic traits are expected to be more similar among closely related species than among species that diverged long ago (all else being equal). This pattern, known as phylogenetic niche conservatism, also applies to traits that are important to determine the niche of species. To test this hypothesis on ecological niches, we analyzed isotopic data from 254 museum study skins from 12 of the 16 species of the bird genus *Cinclodes* and measured stable isotope ratios for four different elements: Carbon, Nitrogen, Hydrogen and Oxygen. We find that all traits, measured individually, or as a composite measurement, lack any phylogenetic signal, which in turn suggests a high level of lability in ecological niches. We compared these metrics to the measurements of morphological traits in the same genus and found that isotopic niches are uniquely evolutionarily labile compared to other traits. Our results suggest that, in *Cinclodes*, the realized niche evolves much faster than expected by the constraints of phylogenetic history and poses the question of whether this is a general pattern across the tree of life.

## Introduction

Phylogenetic niche conservatism (PNC) is the tendency for species to be more ecologically similar to their ancestors and closely related species than expected by chance, even in the face of ecological heterogeneity [1]. Though a certain degree of PNC is an inexorable consequence of descent with modification, as species are likely to inherit ecological features of their ancestors, certain traits show stasis in the rate of change they undergo across phylogenetic trees [2]. PNC has been proposed to explain latitudinal diversity gradients [3–5], and to be of importance for speciation [6,7]. For that reason, PNC has garnered attention over the last two decades and has been the focus of intense research [1, reviewed in 8,9]. In turn, multiple evolutionary processes, including gene flow between lineages [10] and genetic constraints [9,11,12], have been invoked to explain patterns of PNC, which indicates that the study of PNC can be critical to understanding the mechanisms of phenotypic evolution [9]. Examples of phylogenetic niche conservatism in particular traits abound across multiple taxa including birds [13,14], mammals [15,16], frogs [17–19], insects [2,5,20], and plants [4,21–24]. Current tests of niche conservatism often examine the degree to which traits follow the phylogenetic history of a given group [25,26], but usually lack a comparative framework. In spite of the critical importance of PNC as a process to explain genetic divergence, key aspects of the phenomenon remain largely understudied [reviewed in 10]. Paramount among these questions is what organismal traits tend to be conserved as divergence occurs vs. which ones diverge rapidly [10,21]. To date, few attempts have measured the influence of phylogenetic history across a multitude of traits in the same clade, and compared whether there are notable differences among trait classes.

The birds in the genus *Cinclodes* (Furnariidae) are a rapidly diverging [27] lineage of South American suboscine songbirds that have diversified in morphology [28], as well as diet [29–32], migratory behavior [30,32], and residential elevation [32–34]. The genus includes 16 species distributed across South America [34–36]. *Cinclodes*, as well as the remainder of the Furnariidae, may represent a case of continental adaptive radiation [28,37–39] in the last five million years (Figure 1 in Derryberry et al. 2011). The rapid diversification and the exquisite level of phenotypic characterization for the majority of species in the genus offers an excellent opportunity to study whether different traits undergo correlated evolution.

**FIGURE 1.**
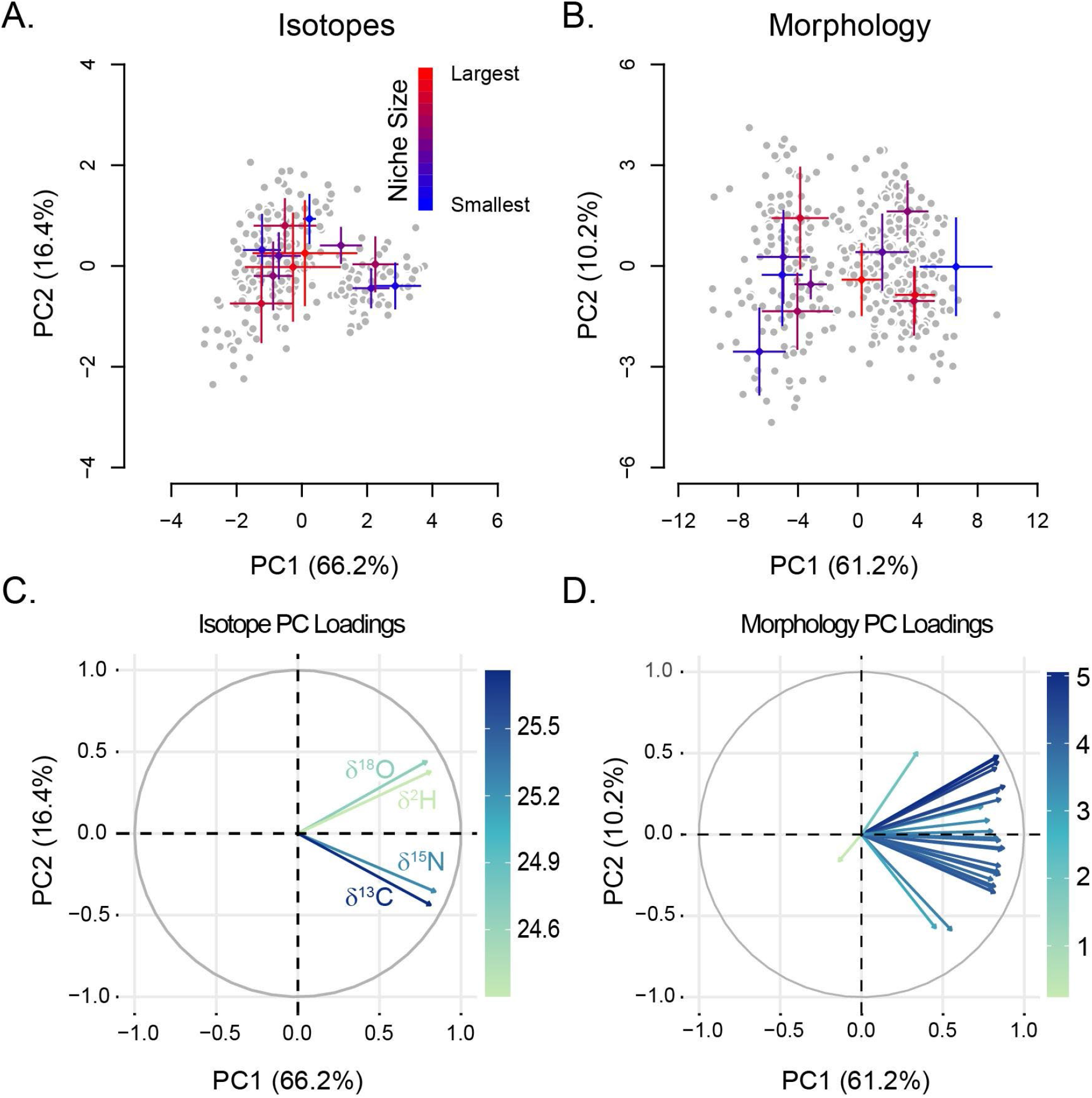
Principal components results for stable isotope values and morphology in Cinclodes. *Cinclodes* isotopic niches measured in PC space (A.). Colored points represent the mean PC1 and PC2 values for each species, with standard deviation error bars. Mean points and error bars are color coded from largest area (red) to smallest (blue). PC1 accounted for 66.2% of the variance, while PC2 captured 16.4%. Pane B. shows the PC1 and PC2 morphospace. Colored points are species mean values and error bars are standard deviations. Mean point and error bar colors follow Pane A. All four isotopic axes loaded strongly on PC1_ISO_ (pane C.), but PC2_ISO_ showed a separation in loadings between the dietary niche axes (C and N) and the geographic niche axes (O and H). All but one of the morphological variables loaded strongly and positively on PC1_MORPH_ (Pane D.); the only exception was wingtip convexity, which had a weak negative loading.

One of the phenotypic axes that has been previously characterized in *Cinclodes* is interspecific variation among isotopic niches. Isotopes of carbon and nitrogen capture variation in trophic gradients (Gannes et al. 1998, Post 2002), while isotopes of hydrogen and oxygen are informative to infer elevational niche (Araguás-Araguás et al. 2000, Newsome et al. 2015). Variation in these four elements is a reliable proxy of the set of resources that are used by an organism [32,40–44]. Previous studies [32,42,45] have reported variation in geographic and the dietary niches in *Cinclodes* and have revealed that species with a wide distribution tend to have the widest diets—measured as isotopic niche widths— an observation that lends support for the resource breadth hypothesis [46]. Nonetheless, none of these studies have quantified the phylogenetic signal of isotopic niches nor compared it with that of morphological traits such as body size and wing shape. These comparisons have the potential to inform not only whether isotopic niches show a phylogenetic signal but also if these traits evolve more readily than morphological traits, presenting a chance to compare whether different trait classes follow a pattern of PNC within a clade. The wealth of data from multiple axes of phenotypic evolution in *Cinclodes* provides the precise opportunity to compare the extent of PNC in multiple traits in a single biological system.

In this study, we used data describing the isotopic niches of 12 species of *Cinclodes* using carbon (δ^13^C), nitrogen (δ^15^N), hydrogen (δ^2^H), and oxygen (δ^18^O) as niche axes, and calculated two phylogenetic signal metrics to look for PNC in the lineage. Additionally, we calculated phylogenetic signal metrics for 23 morphological traits. We find low phylogenetic signal in the mean values and the width of isotopic niches of *Cinclodes*, rejecting PNC for these metrics of ecological niche. On the other hand, we find that the majority of morphological traits show a moderate level of phylogenetic signal; traits that serve as a proxy of body size have a strong signal. These results suggest that even though morphological evolution conforms to the expectations of a PNC pattern, isotopic niches in *Cinclodes*, are much more evolutionarily labile.

## Methods

### Data

To compare phylogenetic signal from isotopic niche breadth to that of other traits in *Cinclodes*, we used two previously published datasets that reported phenotypic values for isotopic and morphological traits for 12 species of the genus. We describe the size and provenance of each of these two datasets as follows.

#### Isotopic niche traits

We used the isotopic trait data from Rader et al. [32]. This dataset included isotopes of carbon, nitrogen, oxygen, and hydrogen from the feather samples for 12 *Cinclodes* species. Sample sizes per species ranged from 3 in *C. pabsti* and *C. comechingonus* to 42 in *C. albidiventris* (Table S1). Isotopes of carbon and nitrogen differ along trophic gradients [47,48], while isotopes of hydrogen and oxygen can capture elevation [41,49]. Isotopic ratios are expressed as δ values:

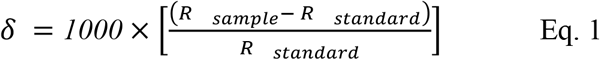

where R_sample_ and R_standard_ are the ratios of ^2^H/^1^H, ^18^O/^16^O, ^13^C/^12^C, or ^15^N/^14^N of the sample and standard, respectively. Isotopic ratios are presented relative to the internationally accepted standards: Vienna Standard Mean Ocean Water (VSMOW, isotopes of oxygen and hydrogen), Vienna Pee Dee Belemnite (VPDB, carbon isotopes), and atmospheric nitrogen (VAIR, nitrogen isotopes). The units are expressed as parts per thousand, or per mil (‰). Precision for δ^2^H was determined by analysis of two exchangeable (keratin) and three non-exchangeable (stearic acid and two oils) organic reference materials, which ranged in δ^2^H from −235‰ to −30‰. Keratin is a good proxy for correction of exchangeable hydrogen [50,51]. Variation (SD) within and among mass spectrometer runs of the five reference materials used for δ^2^H was ≤4‰. All samples for δ^2^H analysis were run in duplicate, and the mean difference between duplicates was typically equal to or less than analytical precision. Precision for δ^13^C and δ^15^? was determined by analysis of acetanilide and alanine standards; within and among run variation (SD) was ≤0.2‰ for both δ^13^C and δ^15^N.

#### Morphological traits

We used a subset of the morphological trait data from Rader et al. (2015). This dataset included 23 morphological measurements for 12 *Cinclodes* species (Table S2). Details of how each trait was measured are provided in Rader et al. (2015). Briefly, the dataset includes morphological measurements [as proposed in 52], taken from study skins housed at the American Museum of Natural History (AMNH), the Field Museum of Natural History (FMNH), the Louisiana State University Museum of Natural History (LSUMZ), and the National Museum of Natural History at the Smithsonian Institution (USNM). We included measurements of head and body length (both proxies for body size), bill length, bill width and depth were measured at two points along the bill (at the proximal end of the bill and its midpoint). We also included the lengths of each of the primary feathers along with the central and lateral tail feathers and two wing shape indices [pointedness and convexity;, 53]. Finally, we included measurements of the hindlimbs: tarsometatarsus length, and the lengths of each digit and its claw. Measurements were all taken with digital calipers (Mitutoyo, Model CD-600CX, with a precision of 0.5 mm) interfaced with Microsoft Excel.

### Estimates of isotopic niche and morphospace width

We measured the breadth of isotopic space occupied by each species as a proxy of their isotopic niche widths. We used principal components analysis using the R function ‘*prcomp’* [library *stats*, 54] to reduce the dimensionality of the δ^13^C, δ^15^N, δ^18^O, and δ^2^H isotopic data set into fewer, and orthogonal niche axes. We adopted a Bayesian approach that was developed for other isotopic niche studies [55,56] to estimate the area of standard ellipses (SEA) occupied by each species in PC space as metrics of their isotopic niche width. These areas and their bootstrapped 95% confidence intervals (n=10,000) were estimated, correcting for small sample size, with the ‘*siber*.*ellipses’* R function [library *siar*;,55–57]. We followed a similar approach to reduce the dimensionality of the morphological dataset.

### Phylogenetic Signal

Next, we studied whether the variation in isotopic and morphological traits followed the expectations of PNC by measuring the phylogenetic signal of each trait. We used the *Cinclodes* ultrametric phylogenetic tree inferred by Derrybery et al. [38] which is based on three nuclear (*BF7, RAG1*, and *RAG2*) and three mitochondrial markers (*COII, ND2*, and *ND3*). We pruned the phylogenetic tree for the family Furnariidae, including associated estimated branch lengths and divergence times [38] to include only the *Cinclodes* species for which we had phenotypic data samples using the *‘drop*.*tip’* function in [library *ape*, 58]. The inferred relationships between *Cinclodes* species are based on information from *BF7, COII, ND2*, and *ND3* [Table S1 in 38].

We used this phylogenetic tree to measure phylogenetic signal in niche width using two different, and complementary, metrics: Blomberg’s *K* [59], and Pagel’s *λ* [60]. Blomberg’s *K* (Blomberg et al. 2003) compares the similarity among relatives compared with that expected from a Brownian motion model of evolutionary change along branches of the same length and determines whether the trait value changes randomly, in both direction and magnitude, along the tree (Blomberg et al. 2003). *K* ranges between 0 and ∞. When *K* equals 1, there is a close correspondence between the phenotypic values observed and those predicted by a Brownian motion model. When *K* is lower than 1, phylogenetic relatives are less similar to each other than expected under Brownian motion evolution, and is evidence against a PNC pattern [1,10,15]. If *K* is larger than 1, species are more similar than expected under Brownian motion evolution, which may suggest PNC, though it may not be sufficient to confirm PNC [see 10].

Second, we used the maximum likelihood optimized [61] value of Pagel’s *λ* [60]. *λ* is a proxy of the extent to which the phylogenetic history of a clade is predictive of the trait distribution at the tree tips. The *λ* statistic ranges between 0 and 1; when *λ* is 0 the phylogenetic structure is the equivalent of a star phylogeny, with all tips emerging from a single node and with equal branch lengths. When *λ* is 1, the phylogenetic structure conforms to the Brownian motion expectation as determined by tree topology and branch lengths. To calculate both metrics, we used the ‘*phylosig*’ function in the *Phytools* R package [62] including 1,000 simulations to determine if the calculated value differed from zero. We tested the significance of the two phylogenetic signal metrics by comparing the *K* and *λ* values recovered for the phylogeny to those for an iterative tip-shuffling randomization [61]. For morphological traits, we followed the same approach we used for the isotopic data to calculate phylogenetic signal (See immediately above). Since phylogenetic signal can be a poor predictor of PNC in some instances in which traits evolve following a Ornstein Uhlenbeck Model [OU, 26], we fit six different models of phenotypic evolution (white noise, Brownian motion, Ornstein-Uhlenbeck, Kappa, Early Burst, and Delta) and studied which one best-fit the data. We used the function *fitContinous* in the *geiger* R package [63,64]. To determine whether the phylogenetic signal differed systematically between isotopic and morphological traits, we used a Kruskal-Wallis test [*‘kruskal*.*test’* function in the *stats* R package;,54].

## Results

First, we characterized the extent of phenotypic variation in isotopic niches in *Cinclodes*. Since the isotopic niche is a multitrait syndrome, we reduced the dimensionality of the dataset using PCA. Our analysis of the isotopic niche data found that, as expected, there were interspecific differences in both the position (mean isotope values) as well as the breadth (dispersion of isotope values) on each of the four isotopic niche axes (Figure S1). The migrant species *C. oustaleti* and *C. patagonicus* had broad ranges of values on each of the four isotopic axes, which contrast with the narrow isotopic ranges of the marine specialists *C. nigrofumosus* and *C. taczanowskii* [Figure 1; also see 32]. Terrestrial species with broad elevational ranges (e.g., *C. albiventris, C. albidiventris*, and *C. excelsior*) showed broad ranges on the oxygen and hydrogen isotopic axes, but comparatively restricted ranges of carbon and nitrogen (Figure S1). *Cinclodes antarcticus*, the sole island resident among the genus, had a broad range of carbon and nitrogen values, but relatively reduced variation in oxygen and hydrogen (Figure S1).

The first two PCs explained a cumulative 82.6% of the overall variance (Figure 1A, Figure S1). The values of δ^13^C, δ^15^N, δ^2^H, and δ^18^O all load strongly (i.e., |loading| > 0.3) and positively on the first principal component (PC1), which accounted for 66.2% of the variance (Figure 1C, Figure S2). The second principal component (PC2) accounted for 16.4% of the variance (Figure 1, Figure S2), with δ^2^H and δ^18^O loading positively on the axis and δ^13^C and δ^15^N loading negatively (Figure 1C,D). The estimated standard ellipse areas (SEAs; Figure 2, Figure S3) occupied by *Cinclodes* species in PC space differed in both position (their centroid in PC space, Figure S3) and size (Figure 2, Figure S3A) confirming that, as a whole, isotopic niches differ among *Cinclodes* species. SEAs ranged from 0.64 and 0.67 for *Cinclodes nigrofumosus* and *taczanowskii*, both marine specialists, to 5.11 for *C. patagonicus* and 4.92 for *C. oustaleti*, both migrants (Figure 2). The SEA for *C. comechingonus* was also quite small (0.37), however with a sample size of 3 for this species, we hesitate to interpret this result further. Marine specialists (*C. nigrofumosus, C. taczanowskii*) had positive values of PC1, contrasting with the negative PC1 values of the strictly terrestrial species (Figure 2). The SEAs of the migrant species *C. oustaleti* and *C. patagonicus* had centroids near 0 on the PC1 axis, and spanned the region between the marine and terrestrial specialists. *Cinclodes pabsti* was a notable exception to this observed pattern, as it shows positive PC1 values, despite that it has never been observed feeding on marine resources.

**FIGURE 2.**
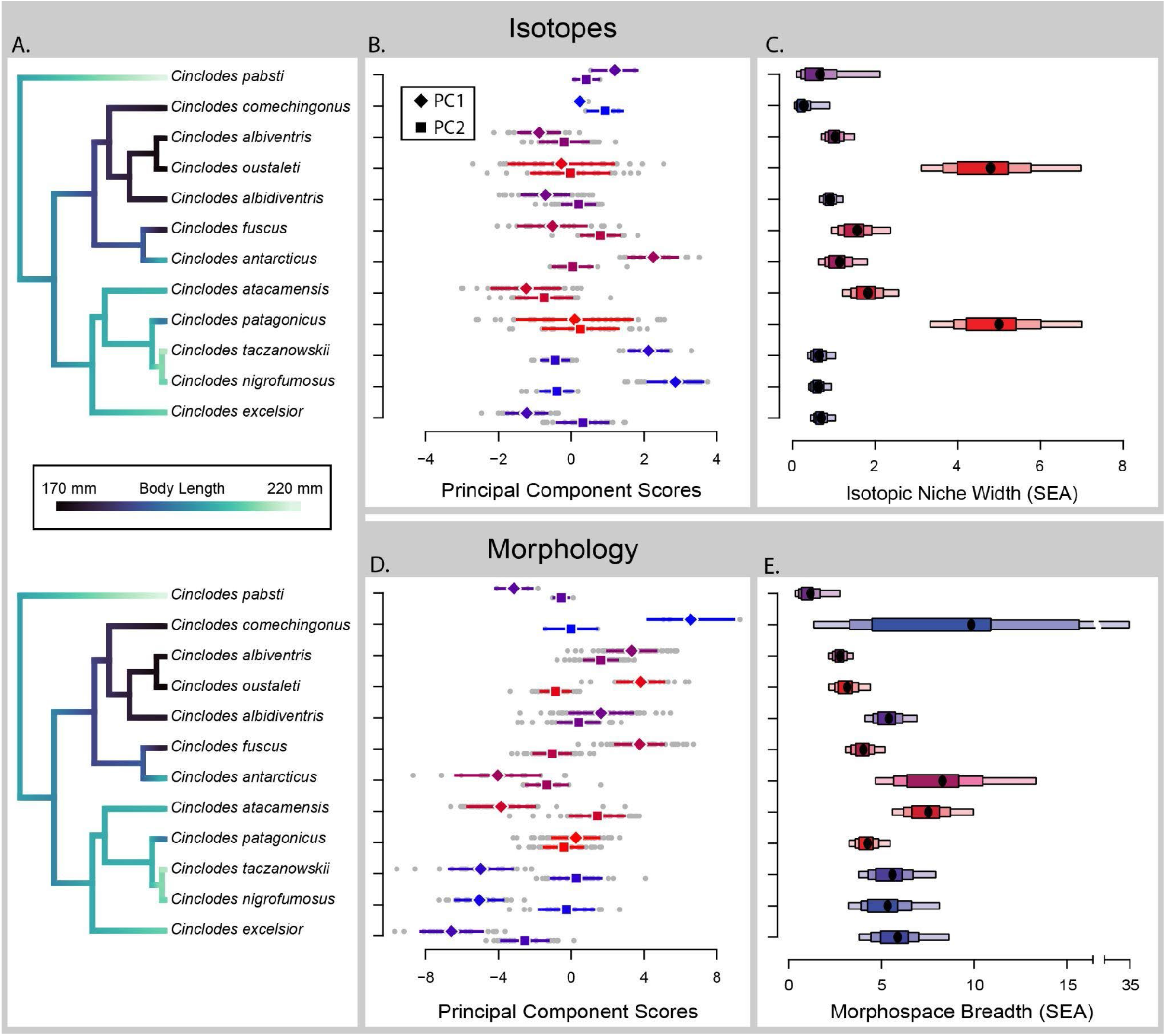
Phylogenetic distribution of isotopic niche widths and morphospace breadth in *Cinclodes*. The *Cinclodes* phylogeny, pruned from the larger furnariid phylogeny from Derryberry et al. [38] is shown in Pane A., with branch length color showing phylogenetic variation in body length. There are two primary clades of Cinclodes, with members of one clade being markedly smaller in body size. The distribution of isotopic PC scores for PC1 (diamonds) and PC2 are plotted at the tree tips in Pane B. Colors as in Figure 1, with red showing the largest isotopic niches and blue the smallest. Pane C. shows density plot of 10,000 Bayesian posterior draws of isotopic niche widths (SEA_ISO_), Colors follow Pane B and Figure 1. Panes D. and E. show the individual PC axes of *Cinclodes* morphospace, and Bayesian estimates of morphospace (SEA_MORPH_), respectively. Colors and symbols follow Panes B. and C.

Next, we quantified the extent of phylogenetic signal of isotopic niches. We used two complementary metrics of signal: Pagel’s *λ* [60] and Blomberg’s *K* [59]. We found that Pagel’s λ was equal to 0 for both PCs (*λ*_*PC1*_=0.00, *λ*_*PC2*_=0.00; Figure 3). Similarly, we found that Blomberg’s *K* was significantly lower than 1 for both PCs (*K*_*PC1*_=0.31, *K*_*PC2*_=0.47; Figure 3), indicating that the isotopic niches of close relatives are less conserved than expected under a pure model of Brownian motion evolution (1,000 randomizations, *p* < 0.001; Fig. 2). Similarly, isotopic niche width—as measured by the SEA— shows a low phylogenetic signal (*λ*_*SEA*.*iso*_=0.00, *K*_*SEA*.*iso*_=0.143). Consistent with this lack of phylogenetic signal, the best fitting model for both PCs and SEA is a white noise model of phenotypic evolution (Table S3). The lack of phylogenetic signal is not exclusive to the composite measures (PC1 and PC2; Figure 3, Table S4) but also applies to the individual isotopic niche axes (Table S5). Collectively, these results indicate that isotopic niches in *Cinclodes* do not follow a PNC pattern and instead species are more dissimilar than expected by shared phylogenetic history.

**FIGURE 3.**
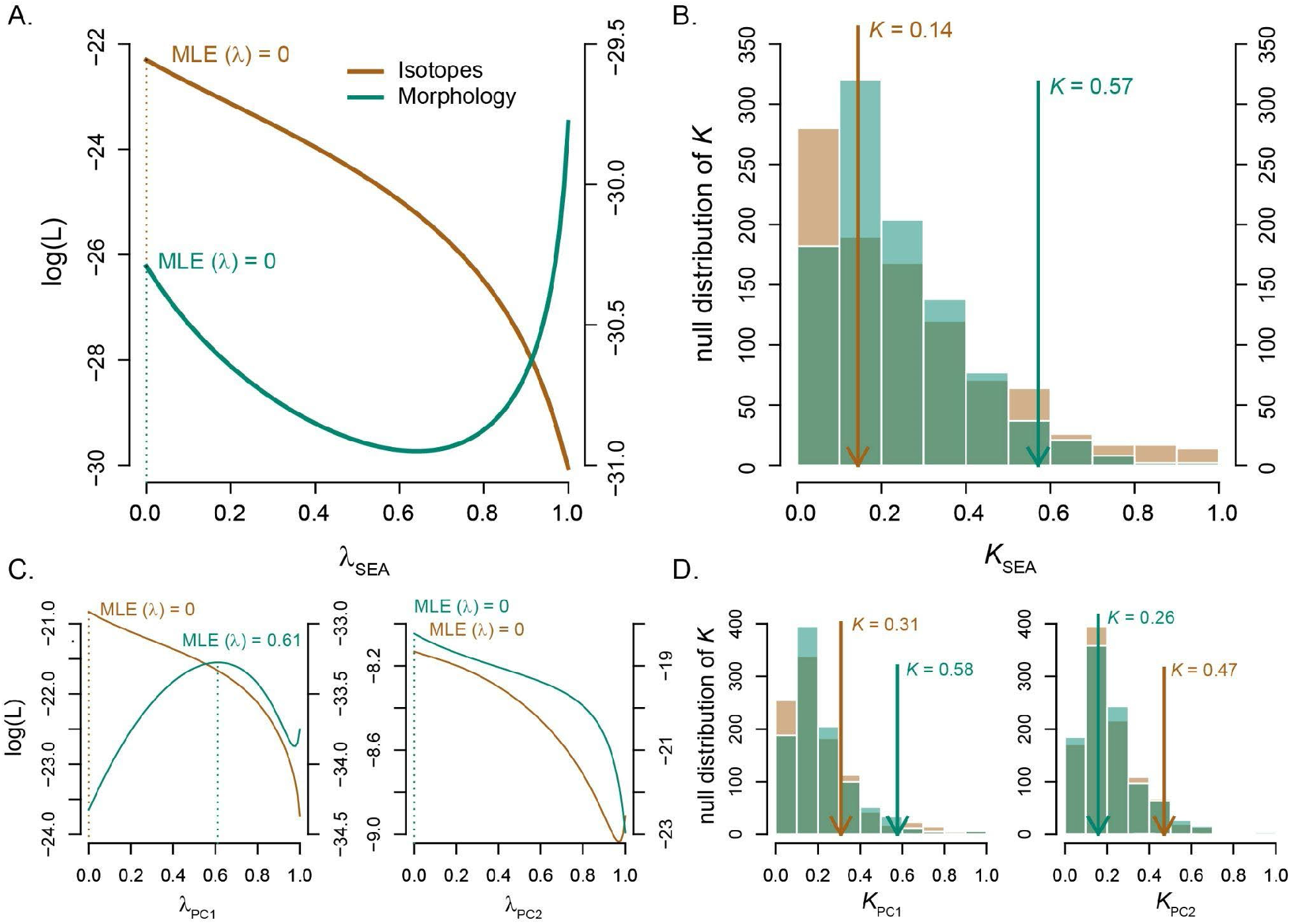
Phylogenetic signal of *Cinclodes* isotopic niches and morphospace. Pane (A.) shows likelihood surfaces of Pagel’s *λ* [60] for isotopic niches (SEA_ISO_; brown) and morphospace (SEA_MORPH_; green). Pane (B.) shows the measured values of Blomberg’s *K* [59] for SEA_ISO_ (brown) and SEA_MORPH_ (green) relative to the results of 1000 tip-shuffling replicates (histograms). Phylogenetic signal is shown in panes C. (*λ*) and D. (*K*) for the first two PC axes in isotopic space and morphospace. There was significant, albeit low, phylogenetic signal for isotopic niches and their constituent PC axes, contrasting with the higher phylogenetic signal for morphological axes.

The lack of phylogenetic signal in isotopic niches poses the question of whether other traits in *Cinclodes* show the same pattern, or whether isotopic niches are somehow unique among phenotypic traits in this clade. To address this question, we compiled previously published data for 25 morphological traits in *Cinclodes* [28] and measured their phylogenetic signal. Similar to the approach we took for isotopic niche, we used a PCA to study morphological traits as a collective phenotype. PC1 explains the vast majority of the variance (61.2%, Figure 1B). As nearly all traits loaded positively on this axis, we interpret it as an axis of body size (Figure 1D). The next three PCs explain ∼20.9% of the variance, cumulatively (Figure S2). This result is largely consistent with previous analyses of morphometry in *Cinclodes* [28]. We calculated the phylogenetic signal of the first four PCs (PC1-PC4). PC1 shows an intermediate value of (*λ*_*PC1*_ = 0.61) which differs both from 1 and from 0 (Figure 3C, Table S3); similarly Blomberg’s *K* (*K*_*PC1*_=0.58) showed a value lower than 1 but significantly larger than zero (Figure 3D, Table S3). The other three minor PCs show different patterns. PC2 shows low values for both metrics (*λ*_*PC2*_ =0.00, *K*_*PC2*_=0.26) both of which differ significantly from 1 and suggest little to no phylogenetic signal as determined by either metric. On the other hand, PC3, and PC4 both show strong phylogenetic signals (*λ*_*PC3*_ =1.00, *K*_*PC3*_=0.90; *λ*_*PC4*_ =1.00, *K*_*PC4*_=1.15). PC1, PC3, and PC4 evolve following a Brownian model of phenotypic evolution, while PC2 evolves according to a white-noise model (Table S3).

Twenty-two individual morphological traits showed a phylogenetic signal significantly higher than zero, and only three showed a *λ* equal to zero (Figure 4). Of the morphological traits, head length and body length are of note because they are proxies of body size [28]. We also compared the values of *λ* and *K* in individual isotopic traits and individual morphological traits (Kruskal-Wallace test, *p* < 0.001 for both cases; Figure 4). We find that out of the 25 individual morphological traits, 22 traits show higher levels of signal (in both *λ* and *K*, Figure 4, Tables S4 and S6) than the isotopic traits (Tables S4, S5). These comparisons suggest that as individual traits, and not only as composites, morphological traits also show a higher level of phylogenetic signal than isotopic traits (Figure 4). SEA’s in PC morphospace ranged from 1.36 (*C. pabsti*) to 8.68 (*C. antarcticus*). Blomberg’s *K* of morphospace SEA’s was moderate (K_SEA.morph_ = 0.57; Figure 3), however the maximum-likelihood estimation of *λ* produced a likelihood surface with two local maxima, one at *λ* = 0 and one at *λ*= 1 (Figure 3). Because the *λ* parameter is dropped from the model when *λ* = 0, the MLE optimized value of *λ*_SEA.morph_ = 0. The difference between the likelihood values is low (−30.22 at *λ* = 0 vs. −29.67 at *λ* = 1), and *λ*_SEA.morph_ cannot be statistically assigned to either maximum.

**FIGURE 4.**
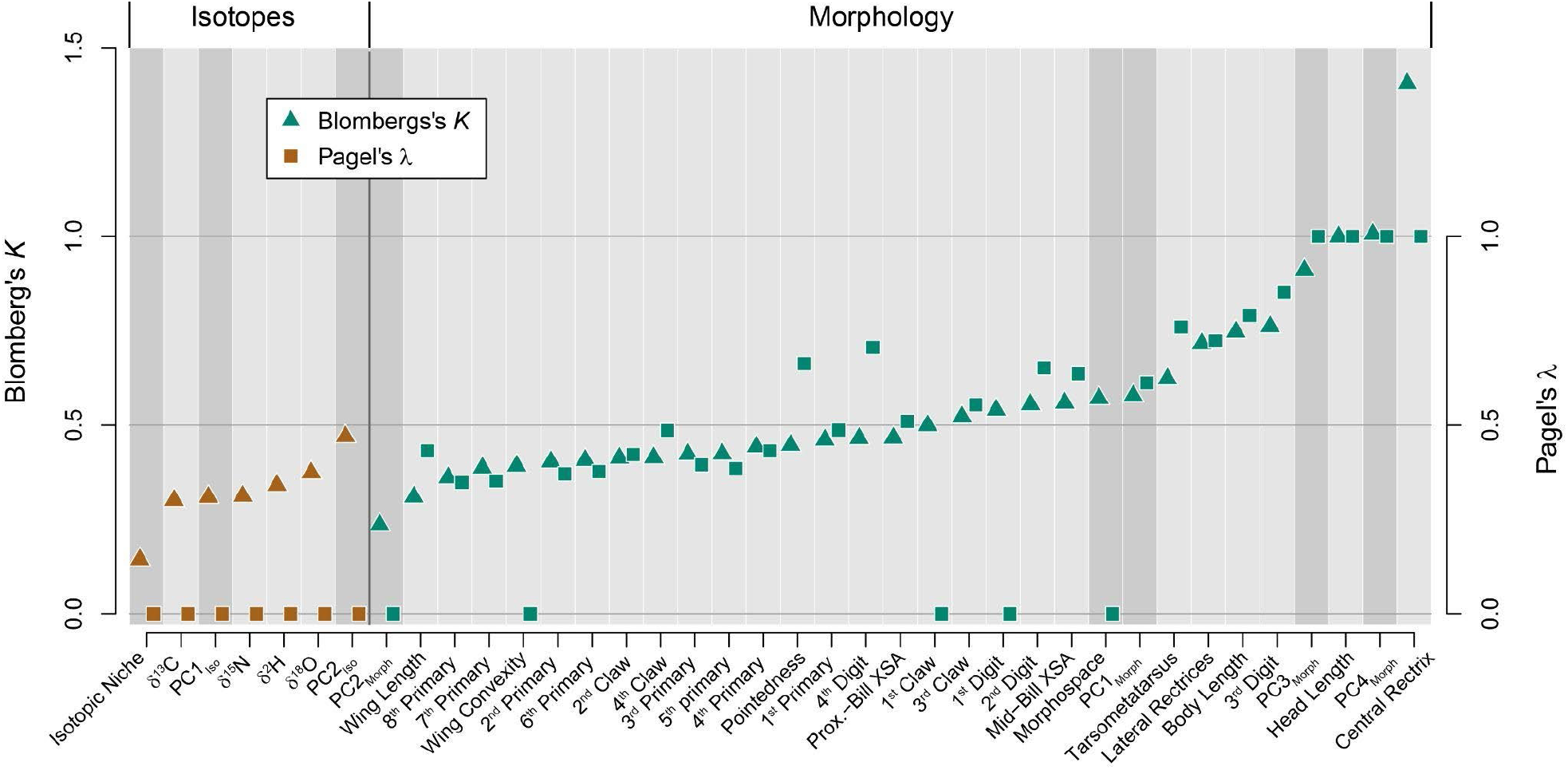
Comparison of phylogenetic signal between isotopic and morphological datasets. Phylogenetic signal for each trait in our study, ranked from lowest to highest *K* (square markers). Triangle markers show Pagel’s *λ*. Isotopic traits are denoted by brown markers, while green represents morphological traits. Individual traits are set against light gray backgrounds and composite traits (PC axes and SEA’s) against darker gray backgrounds.

## Discussion

The study of niche conservatism and divergence is a fundamental component of modern ecology and represents the opportunity to understand how functional diversity is assembled over time. Stable isotope analyses are a scalable approach to study the ecological niche of the species within a clade and thus allow for comparative analyses [32,41,65,66]. The isotopic niche of each species can be viewed as a proxy of the breadth of two fundamental ecological axes: trophic gradients [47,48], and elevational niche [41,49]. Previous work in the bird genus *Cinclodes* demonstrated that species in this genus differed in their isotopic niche widths [32]; trophic niche widths increased with species’ geographic niche widths, supporting the resource breadth hypothesis [46]. We used the same dataset to study how isotopic niches have changed over time in *Cinclodes*. We find that isotopic niches in this bird genus have little to no phylogenetic signal. This absence of phylogenetic signal might suggest that the rate of evolution in the traits that determine realized niche in this bird genus evolved fast enough to not be constrained by phylogenetic relationships. This in turn constitutes evidence against the hypothesis that isotopic niches follow a pattern of PNC in *Cinclodes*. In contrast, morphological traits in the same genus tend to have strong phylogenetic signal, suggesting that at least some traits are indeed constrained by phylogeny in *Cinclodes*. Our results are consistent with the idea that morphological traits are more affected by phylogenetic history than dietary and physiological traits [67].

The absence of PNC in isotopic niches observed in *Cinclodes—*as evidenced by their low phylogenetic signal—is not universal across life but it is not unique either. In corals, Nitrogen and Carbon isotopes show a strong signal of phylogenetic conservatism [Pagel’s *λ* > 0.7 in all cases, Figure 2 in 68]. Notably, species assemblies from the Caribbean show higher phylogenetic signal than assemblies from Polynesia (*λ*_*Caribbean*_ ≈ 0.9 vs. *λ*_*Poynesia*_ ≈ 0.7), which highlights the existence of differences between locations in the importance of environmental and biotic factors on determining the extent of isotopic niche in corals [68]. Similarly, the extent of phylogenetic signal in Ericaceae plants is variable and depends heavily on environmental conditions [69]. While riparian, swamp, and peatland species show low signal in their composite isotopic niche (*λ* < 0.38); rock, barren, and sandy soil species show strong signal [both *λ* values > 0.77, Table 2 in 69]. On the other hand, akodontic rodents show low levels of phylogenetic signal in both Nitrogen and Carbon [Blomberg’s *K*=0.36 for both traits, 70]. These results emphasize the heterogeneity in the rate and mode of evolution in realized niches and its isotopic proxies, highlighting the need for a systematic comparison of the phylogenetic signal of isotopic niches across taxa using common approaches and metrics.

Our study is not devoid of caveats. First, the sampling of the four isotopes in these studies from museum specimens might be too coarse to provide enough resolution to detect phylogenetic signal. Other studies have found this not to be the case and have indicated that museum specimens are a reliable source of isotopic data [44,71]. Second, seasonal variation in resource use among several species of *Cinclodes* might affect the apparent divergences among species [30]. Both of these limitations would bias our results toward finding phylogenetic conservatism, so they are unlikely to be problematic among *Cinclodes*. Nonetheless, the effect of museum storage and the potential for seasonality are questions that remain unexplored in isotopic niche studies and deserve careful treatment.

Isotopes are not the only proxy of ecological divergence, and other approaches have been used to infer how ecological niches have diverged over time. A rich set of literature has used the species geographic range across clades to infer the climatic niche of species [2,e.g., 4]. Angiosperms and sandflies both show high levels of phylogenetic signal in climatic niche traits which, in turn, supports the hypothesis of tropical niche conservatism for both of these clades [2,4]. Consistent with this hypothesis, divergence events involving transitions across latitudinal bands are rare in both of these clades [2,4]. An integration of geographic occurrence data with that of isotopes has revealed differences between the environmental requirements of a species and the actual realized niche in multiple clades. Carnivoran species, for example, show—as expected—a smaller realized niche compared to the environmental conditions in which they could occur [e.g., 72]. These integrative efforts have also revealed an unprecedented amount of dietary plasticity across species. Similarly, analysis of stable isotopes have revealed that the invasive crayfish *Pacifastacus leniusculus* shows a high level of conservatism in its trophic position (a proxy of the realized niche) across geographic locations in spite of its expanded range and a correlated rapid expansion of climatic tolerance across its invasive range [73]. Further research is needed to assess whether realized and potential niches have different levels of phylogenetic signal, and in turn whether they differ from other morphological traits which have been proposed to be more constrained by phylogenetic history [59,74, but see 75].

## Acknowledgements

We thank Andrius Dagilis, Adam Stuckert, Sean Anderson, Tayte Anspach and Dariel Cortés for their helpful comments on this manuscript. Carlos Martínez del Rio provided invaluable insights and support throughout this project. Funding for this work was provided by NSF grant IOS-44362 to C. Martínez del Rio and NSF DEB-1737752 to D.R. Matute.

---

**FIGURE S1.**
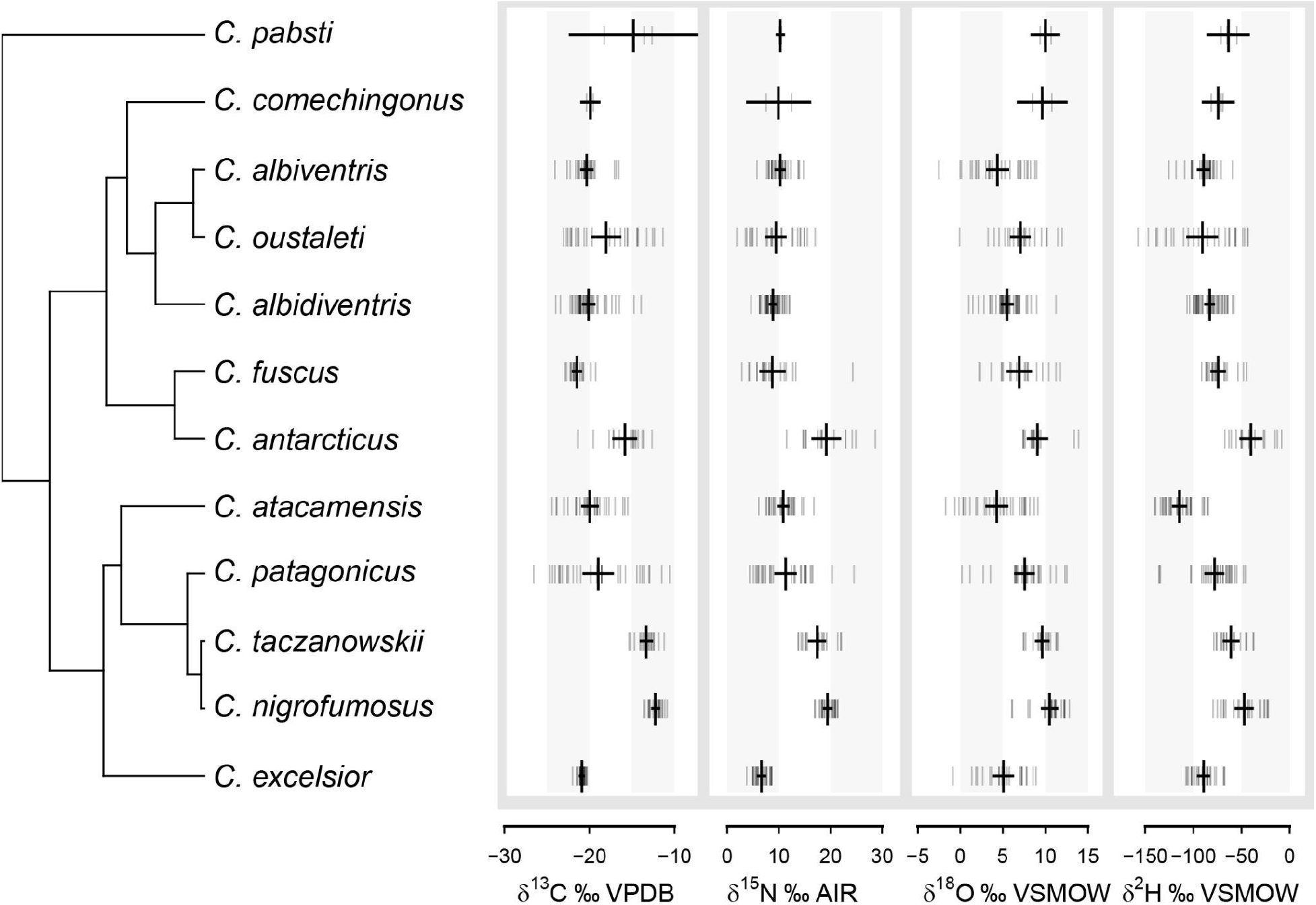
δ^13^C, δ^15^N, δ^18^O, and δ^2^H values of *Cinclodes* species. Light vertical hashmarks are individual data points, and the larger, dark vertical marks are species means. The horizontal bars show the 95% confidence intervals. Migrant species (*C. patagonicus* and *C. oustaleti*) had the broadest ranges on all isotopic axes, while species that were restricted to small geographic ranges (*C. excelsior, C. taczanowskii*, and *C. nigrofumosus*) had the smallest.

**FIGURE S2.**
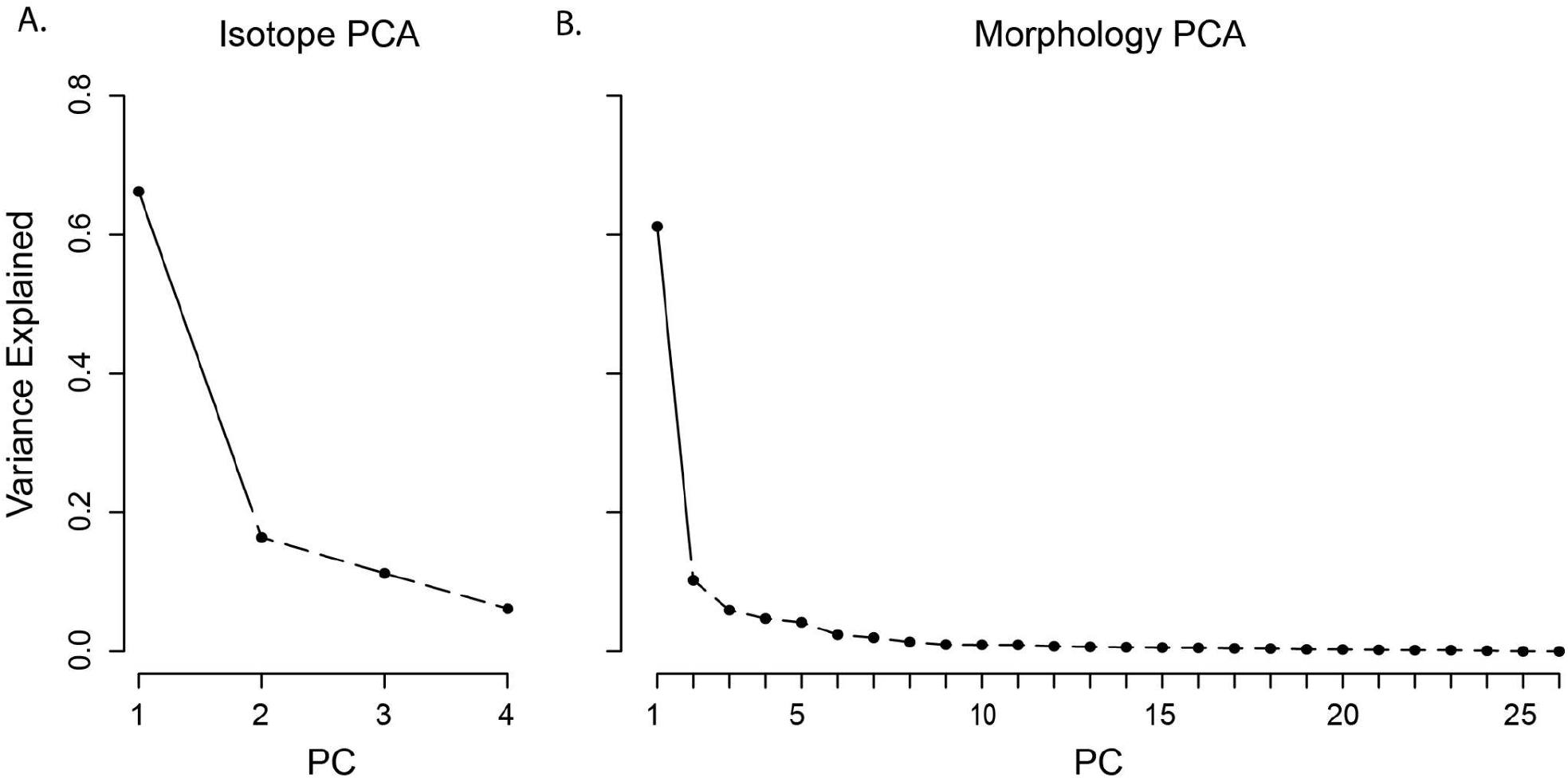
Variance explained by principal component axes in isotopic and morphological datasets. Scree plots showing variance captured by each PC axis in the isotopic (A.) and morphology (B.) PC analyses. Solid vs. dashed lines depict the results of a broken stick analysis: only PC1 from each PCA was significant.

**FIGURE S3.**
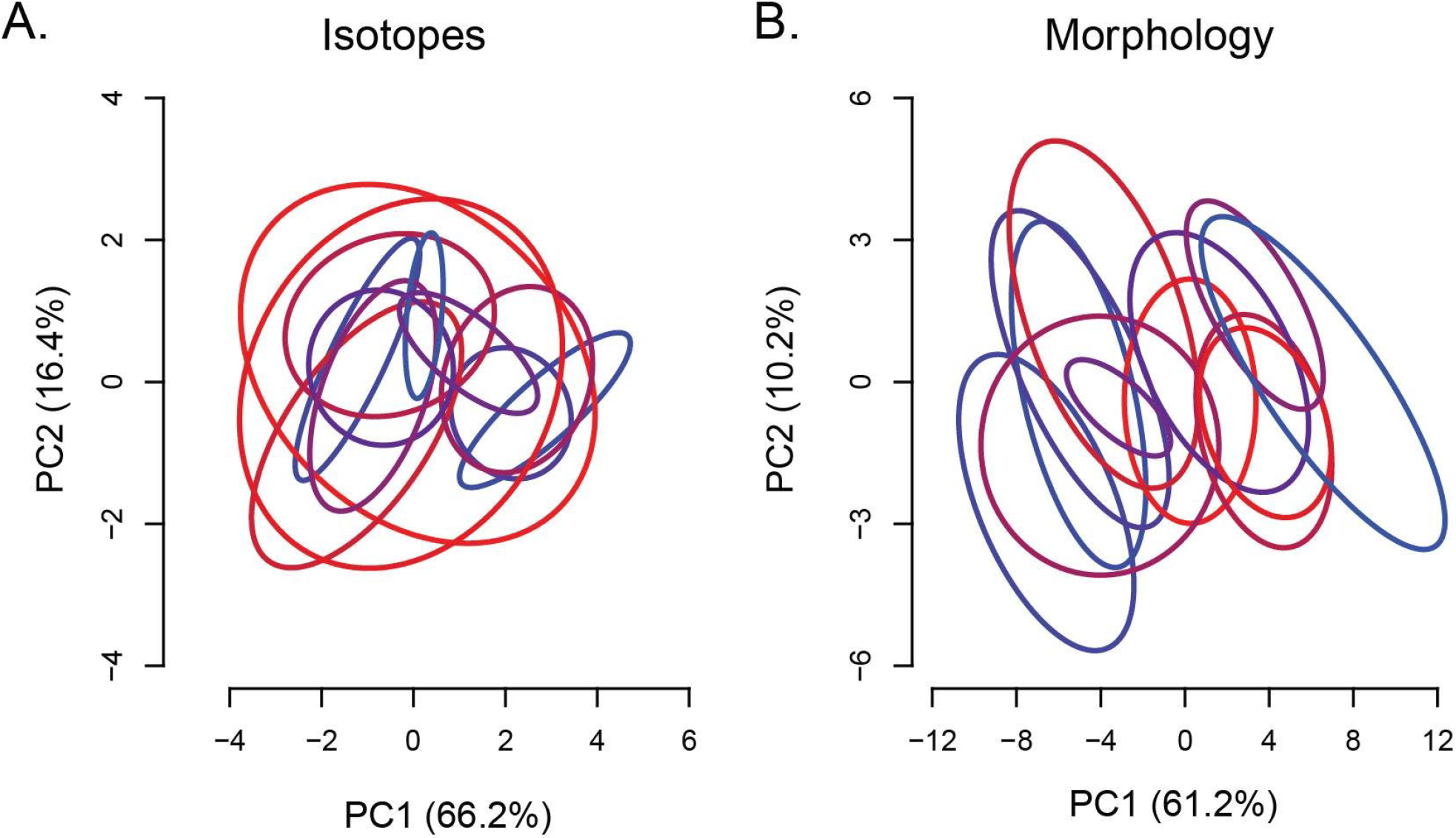
Comparison of isotopic vs. morphological spaces (SEA). Large points are species tips, and smaller black points are internal nodes of the tree. Colors as in Fig. 1, with large isotopic niches in red and small niches in blue.

**FIGURE S4.**
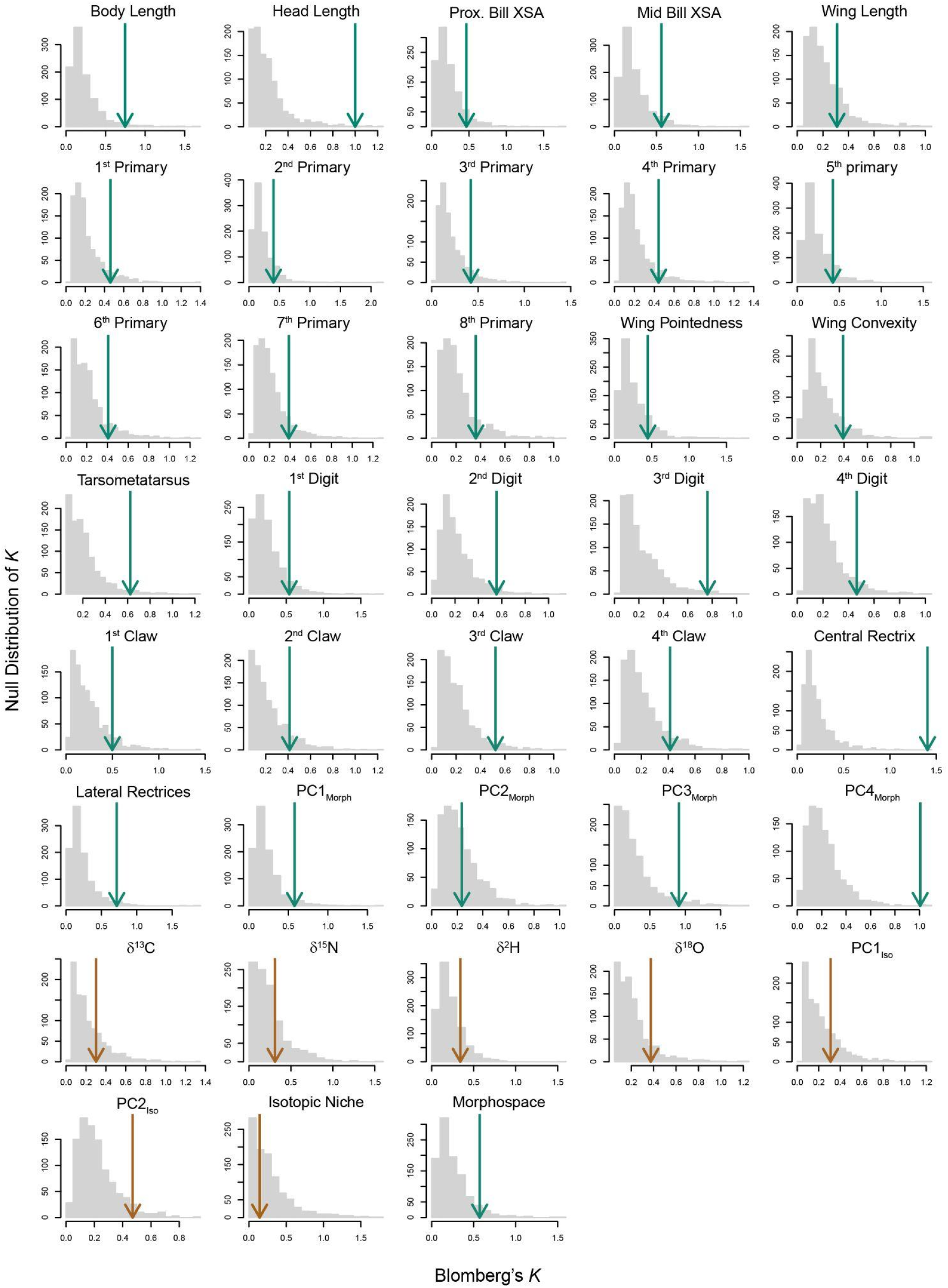
Blomberg’s *K* for each measured trait. Measured *K* for isotopic traits is shown with brown arrows and green arrows show *K* for morphological traits.

**FIGURE S5.**
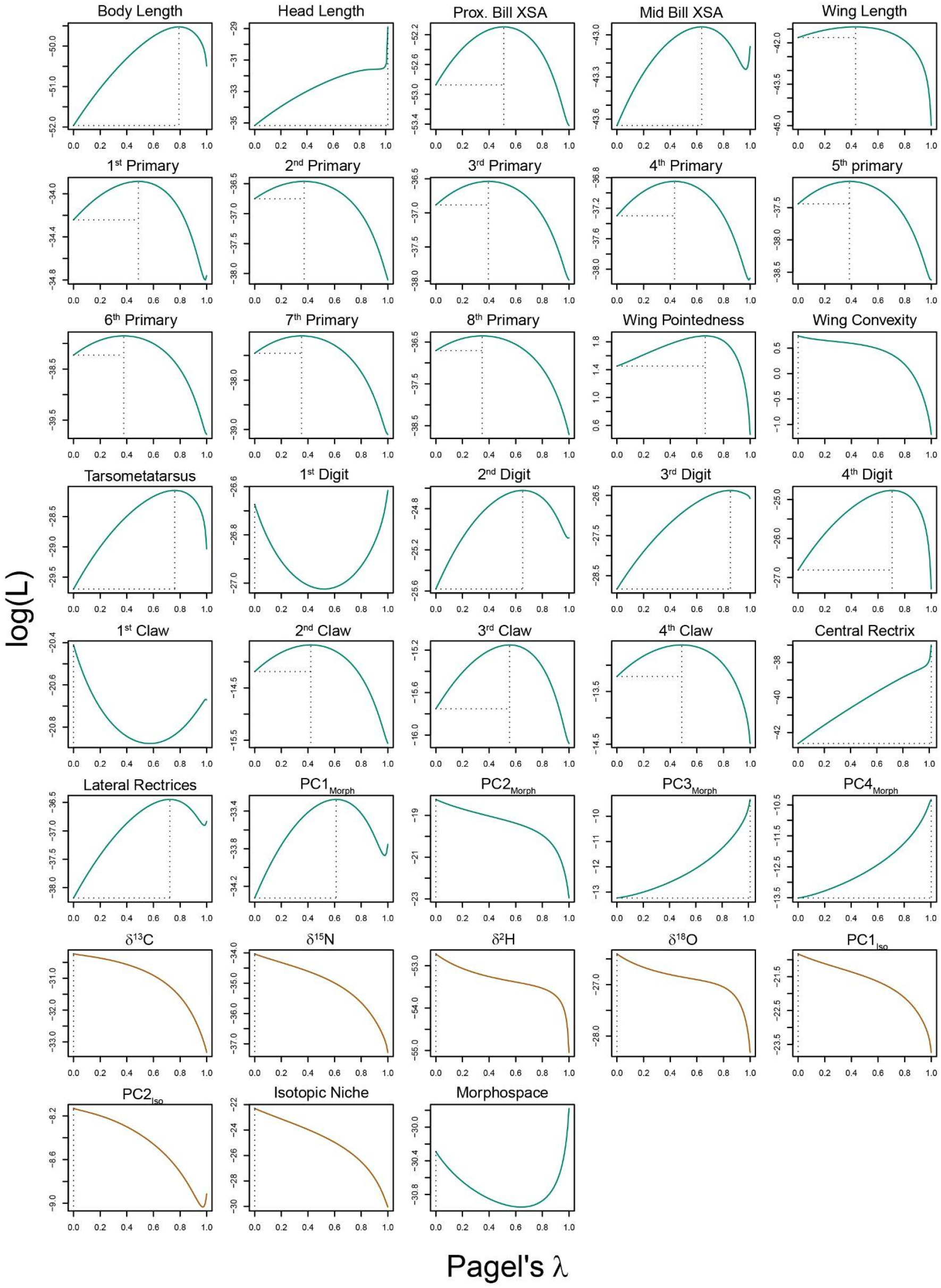
Pagel’s *λ* for each measured trait. Measured *λ* for isotopic traits is shown with brown arrows and green arrows show *λ* for morphological traits.

**TABLE S1.**
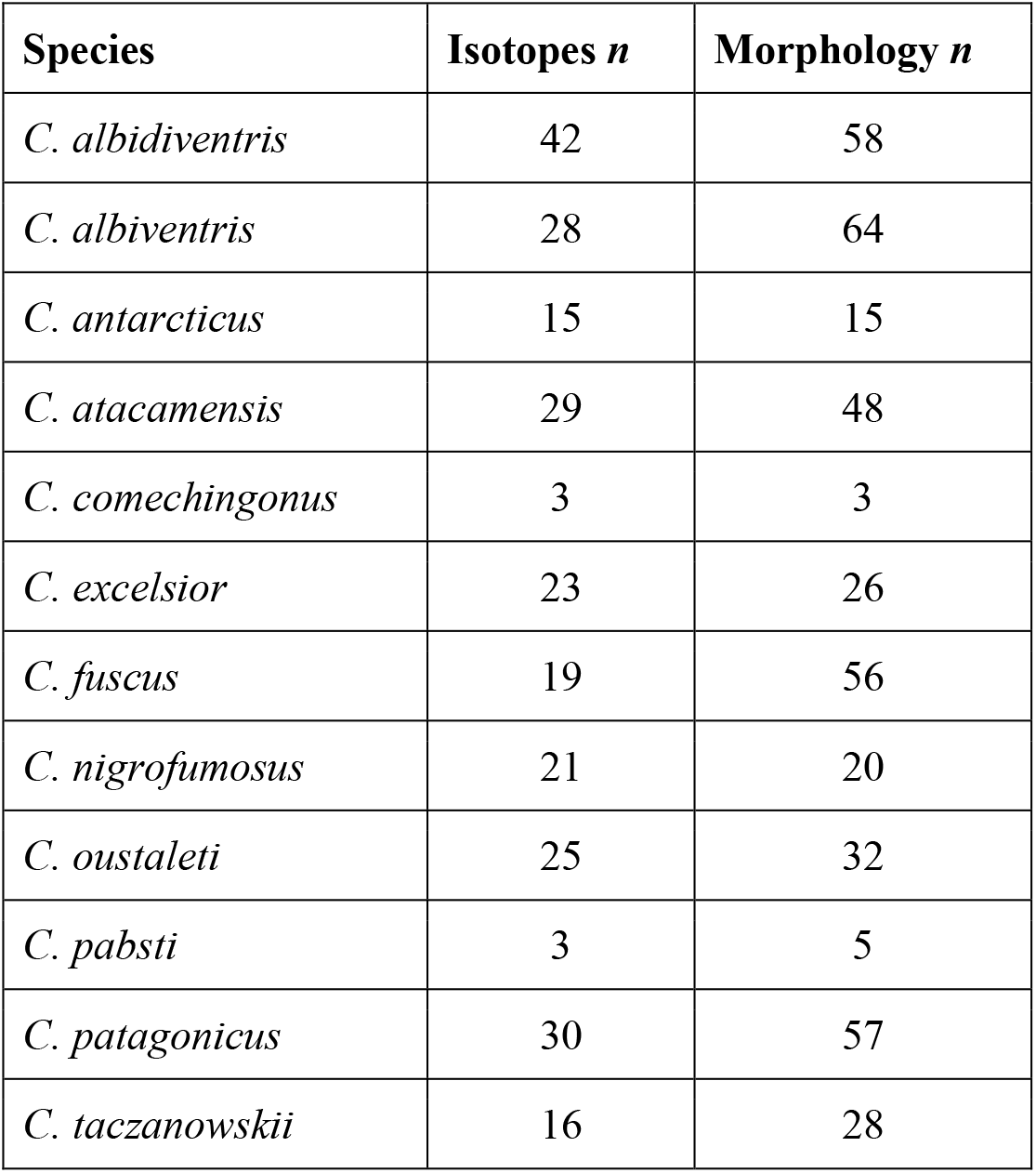
Sample sizes for each species of *Cinclodes*.

**TABLE S2.**
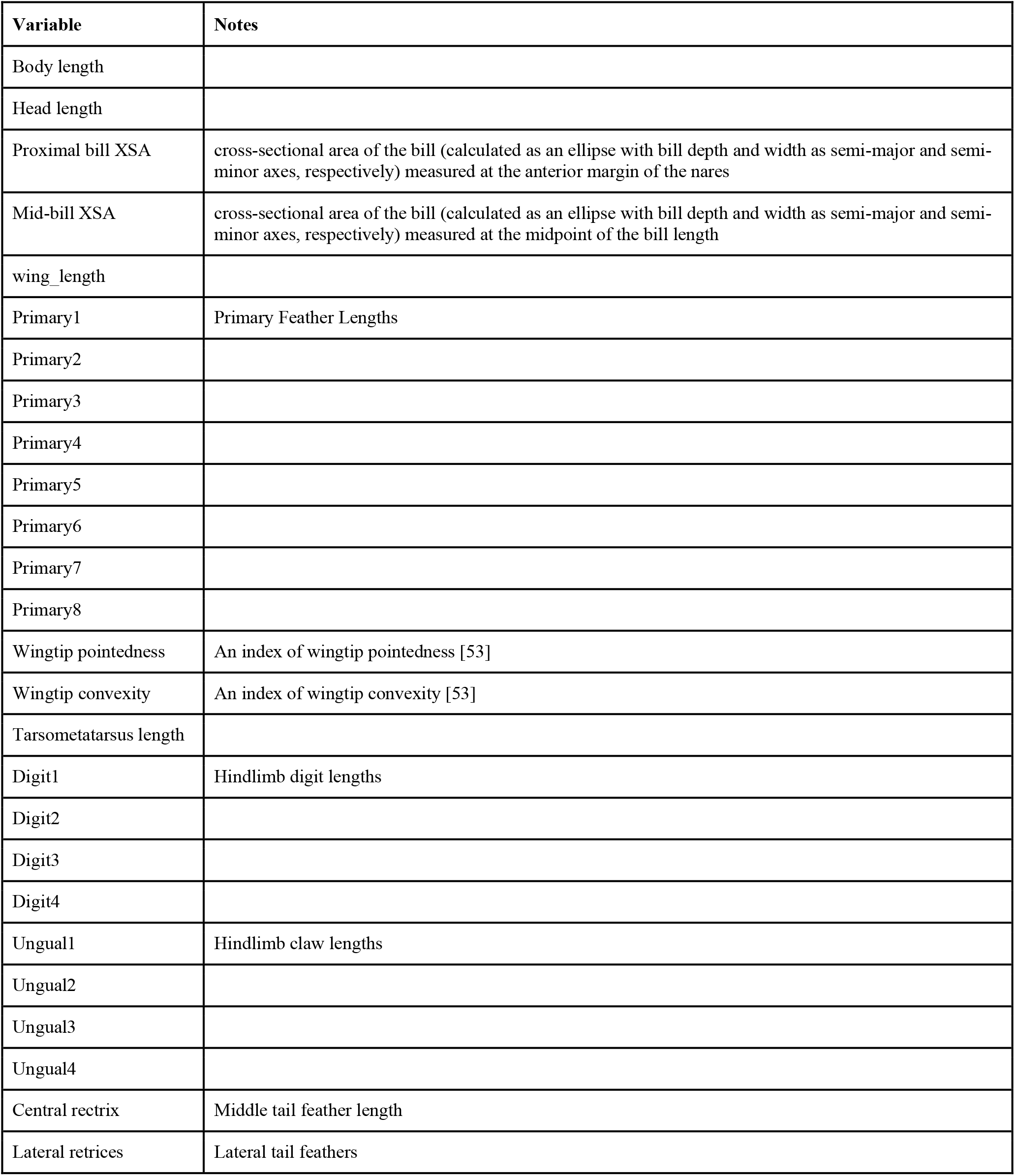
Morphological traits.

| Variable | Notes |
| --- | --- |
| Body length |  |
| Head length |  |
| Proximal bill XSA | cross-sectional area of the bill (calculated as an ellipse with bill depth and width as semi-major and semi-minor axes, respectively) measured at the anterior margin of the nares |
| Mid-bill XSA | cross-sectional area of the bill (calculated as an ellipse with bill depth and width as semi-major and semi-minor axes, respectively) measured at the midpoint of the bill length |
| wing_length |  |
| Primary1 | Primary Feather Lengths |
| Primary2 |  |
| Primary3 |  |
| Primary4 |  |
| Primary5 |  |
| Primary6 |  |
| Primary7 |  |
| Primary8 |  |
| Wingtip pointedness | An index of wingtip pointedness [53] |
| Wingtip convexity | An index of wingtip convexity [53] |
| Tarsometatarsus length |  |
| Digit1 | Hindlimb digit lengths |
| Digit2 |  |
| Digit3 |  |
| Digit4 |  |
| Ungual1 | Hindlimb claw lengths |
| Ungual2 |  |
| Ungual3 |  |
| Ungual4 |  |
| Central rectrix | Middle tail feather length |
| Lateral retrices | Lateral tail feathers |

**TABLE S3.** AICc values of evolutionary models.

| Variable | WN | Lambda | Delta | OU | BM | Kappa | EB |
| --- | --- | --- | --- | --- | --- | --- | --- |
| PC1<sub>iso</sub> | **46.99** | 50.66 | 51.92 | 47.91 | 52.81 | 51.51 | 56.48 |
| PC2<sub>iso</sub> | **21.60** | 25.26 | 23.79 | 23.00 | 23.16 | 26.13 | 26.83 |
| SEA<sub>iso</sub> | **49.91** | 53.58 | 63.36 | 53.37 | 65.40 | 58.94 | 69.07 |
| PC1<sub>morph</sub> | 73.98 | 75.55 | 74.19 | 74.07 | **72.84** | 75.58 | 76.50 |
| PC2<sub>morph</sub> | **41.80** | 45.47 | 50.29 | 45.47 | 51.25 | 50.02 | 54.91 |
| PC3<sub>morph</sub> | 31.81 | 28.34 | 27.53 | 28.05 | **24.68** | 28.32 | 28.34 |
| PC4<sub>morph</sub> | 32.35 | 29.68 | 29.67 | 29.68 | **26.02** | 29.68 | 29.67 |
| SEA<sub>morph</sub> | 65.77 | 68.34 | 66.85 | 67.02 | **64.68** | 68.34 | 68.34 |

**TABLE S4.** Phylogenetic signal of principal components axes for both isotope and morphological data.

| Variable | *K* | *λ* |
| --- | --- | --- |
| PC1<sub>iso</sub> | 0.309 | 0.000 |
| PC2<sub>iso</sub> | 0.470 | 0.000 |
| SEA<sub>iso</sub> | 0.143 | 0.000 |
| PC1<sub>morph</sub> | 0.579 | 0.614 |
| PC2<sub>morph</sub> | 0.258 | 0.000 |
| PC3<sub>morph</sub> | 0.898 | 1.000 |
| PC4<sub>morph</sub> | 1.150 | 1.000 |
| SEA<sub>morph</sub> | 0.575 | 0.000 |

**TABLE S5.** Phylogenetic signal of stable isotope ratios in *Cinclodes* feathers.

| Variable | *K* | *λ* |
| --- | --- | --- |
| δ<sup>13</sup>C | 0.301 | 0.000 |
| δ<sup>15</sup>N | 0.312 | 0.000 |
| δ<sup>2</sup>H | 0.341 | 0.000 |
| δ<sup>18</sup>O | 0.374 | 0.000 |

**TABLE S6.** Phylogenetic signal of morphological variables.

| Variable | *K* | *λ* |
| --- | --- | --- |
| body_length | 0.747 | 0.791 |
| head_length | 0.999 | 1.017 |
| proximal_bill_cross | 0.466 | 0.510 |
| mid_bill_cross | 0.559 | 0.636 |
| wing_length | 0.310 | 0.432 |
| primary1 | 0.461 | 0.487 |
| primary2 | 0.403 | 0.371 |
| primary3 | 0.423 | 0.395 |
| primary4 | 0.443 | 0.432 |
| primary5 | 0.424 | 0.385 |
| primary6 | 0.407 | 0.377 |
| primary7 | 0.387 | 0.352 |
| primary8 | 0.361 | 0.348 |
| pointedness | 0.447 | 0.663 |
| convexity | 0.392 | 0.000 |
| tmt_length | 0.623 | 0.760 |
| digit1 | 0.540 | 0.000 |
| digit2 | 0.555 | 0.652 |
| digit3 | 0.761 | 0.852 |
| digit4 | 0.465 | 0.706 |
| ungual1 | 0.499 | 0.000 |
| ungual2 | 0.413 | 0.422 |
| ungual3 | 0.522 | 0.554 |
| ungual4 | 0.414 | 0.486 |
| central_rectrix | 1.405 | 1.016 |
| lateral_retrices | 0.716 | 0.723 |

